# Characterization of tumour interactions with the immune system in an autochthonous mouse model of glioblastoma

**DOI:** 10.64898/2026.05.13.724869

**Authors:** Margarita Lui, Olivia J Makinson, Meghan L Walsh, Taj J Matthews, John Woulfe, Michele Ardolino, Ian AJ Lorimer

## Abstract

**Background:** Glioblastoma is an aggressive and incurable brain tumor. Clinical trials of immune checkpoint inhibitors showed no clinical benefit in glioblastoma when given after surgery. However, a clinical trial in which PD1 inhibition was given prior to second surgery did show pharmacodynamic evidence for activity. This suggests the possibility that immune checkpoint inhibitors may be more effective in a setting where large tumors are present. Here we have studied immune responses to large tumors in an autochthonous mouse model of glioblastoma.

**Methods:** Glioblastoma was induced by transfection with oncogenic plasmids injected directly into the lateral ventricle of neonatal mice. Immune responses were assessed using a combination of spectral flow cytometry and immunohistochemistry.

**Results:** There was a marked immune response to large tumors, with significant increases in CD4 T cells and dendritic cells. T cell changes occurred primarily at leptomeningeal/perivascular border sites. A large proportion of CD4 T cells expressed PD1 and half of these were regulatory T cells. NK cells were also increased in mice with large tumors, but were predominantly in immature states. The mouse model accurately recapitulates the formation of palisading necroses. These contain apoptotic cells and avidly recruit myeloid cells that are induced to express large amounts of TGFβ.

**Conclusions:** Large glioblastoma tumors generate a border site population of PD1 positive T cells that may explain the pharmacodynamic response in neoadjuvant trials, and a palisading necrosis-driven immunosuppressive mechanism that may explain why responses are insufficient to provide a significant clinical benefit.

**KEY POINTS:** The SB mouse model accurately recapitulates immune features of human glioblastoma

Large tumors induce a significant border site immune response

Palisading necroses in large tumors counter this with a strong immunosuppressive response

**IMPORTANCE OF STUDY:** Immune checkpoint inhibitors have not shown efficacy in glioblastoma when used post-surgery, but do show pharmacodynamic activity when used in patients prior to second surgery (*i.e.* neoadjuvant). This suggest the possibility that immune checkpoint inhibition is more effective when large tumors are present. Using a clinically-relevant autochthonous mouse model, we show here that large tumors induce an immune response that is evident in leptomeningeal border sites. Large tumors in this mouse model also generate palisading necroses, a well-known diagnostic feature in glioblastoma tumors. These palisading necroses generate large amounts of TGFβ, providing a mechanism by which large tumors can suppress border site immune responses. This further supports the concept that palisading necroses are drivers of glioblastoma malignancy and suggests novel strategies to enhance responses to immune checkpoint inhibition in this cancer.

## INTRODUCTION

Glioblastoma is an aggressive and incurable form of brain cancer. Current standard treatments are surgery followed by combined radiation and temozolomide chemotherapy (1). These only give small improvements in survival and there is a clear need for more effective therapeutics. Immune checkpoint inhibitors have been very effective in a number of different cancers; however large, randomized clinical trials of these in glioblastoma did not show any clinical benefit (2–5). These trials were generally done in patients that had undergone previous surgery. Cloughesy *et al*. subsequently performed a small trial in which glioblastoma patients that were undergoing a second surgery were randomized to immune checkpoint inhibitor treatment before and after surgery (neoadjuvant), or treatment after surgery only. Detailed correlative studies showed that there was biomarker evidence for an immune response in the neoadjuvant group, although the data suggested that this was held in check by immunosuppressive macrophages (6–8). There are several possible reasons for the activity in the neoadjuvant setting and a detailed understanding of this may help the development of effective immunotherapy for glioblastoma.

The brain has for many years been viewed as an immunoprivileged site, based on the observation that tissue could be successfully transplanted into the brain but was rejected if transplanted at other sites. This concept is undergoing extensive revision, driven by the recognition there is in fact extensive communication between the brain and immune systems (9). Much of this takes in place in border sites including the meninges, perivascular regions and choroid plexus. These sites are often underrepresented or absent in patient surgical samples. Detailed studies of border site engagement and immunotherapy require immunocompetent animal models that accurately reproduce essential features of glioblastoma in patients (reviewed in (10)). The SB model originally described by Wiesner *et al*., in which glioblastoma is generated *in situ* with relevant oncogenic drivers, may be useful in this regard (11–13). This model induces glioblastoma by intracerebroventricular injection of plasmids in neonatal mice. Plasmids encode either glioblastoma oncogenes, or short hairpin RNAs and constitutively active proteins that mimic the effects of tumour suppressor loss. These sequences are configured as Sleeping Beauty transposons to allow integration, and a plasmid encoding the Sleeping Beauty transposase SB13 is cotransfected to promote integration. As this model generates glioblastoma *in situ*, it avoids possible artifacts due to orthotopic injection of cultured cells and permits the study of early and late disease. It can be configured with different oncogenes (13) and can readily be used in different mouse backgrounds to study host influences on therapeutic responses. This model has been used to study various aspects of glioblastoma (14, 15) and cells derived from this model have been used to study immune checkpoint inhibition (16). Drawbacks of the model are that neonatal intracerebroventricular injections can be challenging and mice develop tumours relatively slowly. Here we describe the use of an improved method for intracerebroventricular injections that makes this technique more accessible and reduces time to tumour formation. We also describe a detailed analysis of immune cell engagement and confirm that the model accurately reproduces the immune cell engagement seen in human glioblastoma. Large tumors in the model actively engage the immune system in the meninges and perivascular regions, inducing increases in T cells at these border sites. Large tumors also develop the palisading necrosis that is diagnostic for human glioblastoma. We show that these regions are a major source of TGFβ that can counter border site immune responses, potentially promoting the regulatory T cell increases and repression of NK cell maturation that are observed in the model.

## METHODS

### Plasmids

PT2/L-Luc//PGK-SB13 (Addgene #20207, 9842 bp); pT2/shp53/GFP4 (Addgene # 20208, 7371 bp); pT/Caggs-NRASV12 (Addgene #20205, 6300 bp); PT3.5/CMV-EGFRVIII (Addgene #20280, 8334 bp) were purified using an endotoxin-free protocol (Endo-free Plasmid Maxi Kit, Qiagen) and identity was confirmed by restriction digests.

### Intracerebroventricular injections in neonatal mice

3D-printed adaptors for a stereotaxic injection frame were made by Polyunity Tech Inc. (Ottawa ON, Canada). An initial version was made following the design described by Olivetti *et al*. (17) and a subsequent modified version was made after testing this. Details of this design are available from the manufacturer. Pregnant FVB/N mice were from Charles River. A detailed protocol for the injections is given in the supplemental information. Plasmid injections were done in the left lateral ventricle. All mouse experiments were done under a protocol approved by the University of Ottawa Animal Care Committee (OHRIe-4326). For survival studies, mice were euthanized at the first signs of morbidity.

### In vivo imaging for luciferase

Optical imaging was done at the University of Ottawa Preclinical imaging core using a Newton 7.0 FT500 imaging system. Mice were injected i.p. with 100 uL of 5 mM CycLuc1 as described (18) and imaged 10-15 mins post injection on open aperture and 12×12 binning sensitivity for 12 minutes.

### Spectral flow cytometry analysis of immune cell populations

Tissue processing and flow cytometry of splenocytes and tumor infiltrating lymphocytes was performed as described (19–21). The left hemisphere of the brain was dissected and dissociated to single cells using the Tumor Dissociation Kit and the gentleMACS Octo Dissociator (Miltenyi Biotec). Cells were homogenized using 40μm cell strainers and red blood cell lysis was performed using ammonium chloride-potassium (ACK) lysing buffer. Spleens were homogenized using 40μm cell strainers. Red blood cell lysis was performed using ACK lysing buffer. Cells were incubated with BD Mouse Fc Block (2.4G2) and then labeled for 30 min at 4°C with the appropriate monoclonal antibodies. Intracellular labeling was performed using Foxp3 fixation/permeabilization buffer (eBioscience) and labeling for 45 min at 4°C with the appropriate monoclonal antibodies. Flow cytometric analysis was performed on an Aurora 5L spectral flow cytometer (Cytek) and data was analyzed with FlowJo 10.10 software.

### Immunohistochemistry

Immunohistochemistry was performed at the Louise Pelletier Core Facility (RRID: SCR_021737) at the Department of Pathology and Laboratory Services, Faculty of Medicine, University of Ottawa. CD3e recombinant rabbit monoclonal antibody (6G6K9) was from ThermoFisher and was used at 1:500; FoxP3 (D608R) rabbit monoclonal antibody was from Cell Signaling Technology and was used at 1:200; Iba1 antibody [EPR16588] (ab178847) rabbit monoclonal antibody was from Abcam and was used at 1:8000; TGF beta-1 Recombinant Rabbit Monoclonal Antibody (PD00-17) was from ThermoFisher and was used at 1:200. Cleaved caspase-3 rabbit monoclonal antibody (cat.# 9664) was from Cell Signaling Technology and was at 1:500. Whole section digital images were generated using a Zeiss Axioscan.Z1 slide scanner and analyzed using Zen2 (blue edition) software.

### Statistics

Graphing and statistical analyses were done using SigmaPlot 14.5. Graphs show mean ± standard deviation unless indicated otherwise. Two-tailed Student t tests were used unless indicated otherwise, after confirming that data were normally distributed. P < 0.05 was considered significant.

## RESULTS

### Tumour take and survival

As described in the original publication, we performed intracerebroventricular injections of plasmids encoding: SB13 transposase and luciferase; EGFRvIII; constitutively active NRAS; short hairpin RNA targeting TP53 and eGFP (11). Similar to the original publication, FVB/N mice formed tumours that were detectable by optical imaging of luciferase activity, and median survival of mice was 85 days, with eight of nine mice developing tumours (Figure 1A). Precise lateral ventricle injection in neonatal mice is challenging with the standard stereotaxic apparatus used for adult mice. Olivetti *et al*. have described an adaptor to facilitate neonatal stereotaxic injections and used this to label neural circuits in developing mice (17). We had their design 3D printed and tested it; we then made a second version with some additional alterations. Using this with the same plasmid amounts and mouse strain as above, the median survival was decreased to 44 days, with all mice developing tumours (Figure 1A). This indicates that, without the adaptor, imprecise injections were limiting transfection efficiency and increasing the time to tumour formation. Use of the adaptor makes this model considerably faster and should make the model easier for new users to adopt. Gross pathology of late stage tumours showed highly invasive disease, with invasion across the corpus callosum into the opposite hemisphere occurring routinely (Figure 1B). Tumours also showed hemorrhagic and necrotic cores, with characteristic palisading necrosis (Figures 1 C and 1D). Early stage tumour formation in the subventricular zone of the injected hemisphere could be detected using the GFP encoded by one of the plasmids (Figure 1E).

**Figure 1:**
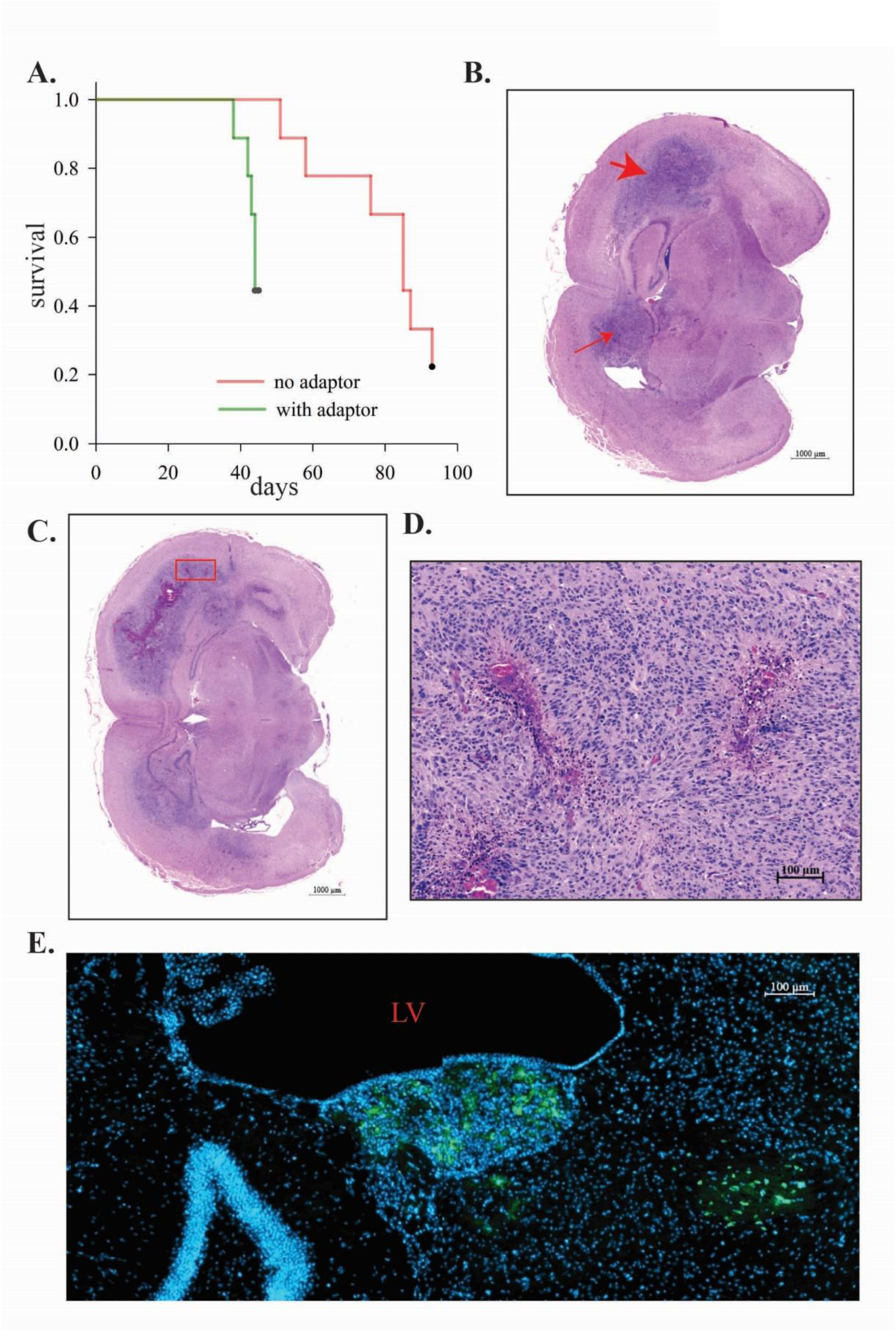
A. Survival without and with stereotaxic surgery adaptor. Neonatal FVB/N mice were injected without using the adaptor (red) or with the adaptor (green) and euthanized at humane endpoint. Median survival times were 85 days and 44 days (P = 0.011 by Log Rank test). For the mice injected without the adaptor, one of two surviving mice showed the presence of tumour after euthanasia; for the mice injected with the adaptor, all remaining mice showed tumours*. B.* Example of late stage tumor. The hemisphere where the lateral ventricle injection was performed is shown at the top (large red arrow). Extensive invasion along the corpus callosum into the opposite hemisphere (small red arrow) is evident. *C. Hemorrhaging and necrosis in tumour at humane endpoint*. A different section from the same mouse as C is shown. *D. Palisading necrosis.* A close up of the section shown in C is shown (area marked by red box). E. Example of an early stage tumor detected by fluorescence microscopy for GFP. LV, lateral ventricle.

### Immune profiling

Immune cell infiltrates in early and late stage tumors were characterized using spectral flow cytometry. Neonatal mice were injected with plasmids and monitored by optical imaging. Samples were collected at a timepoint post injection when only a subset of mice showed tumour formation on imaging; tissue from mice that had detectable tumours by optical imaging is compared with tissue from mice that did not. All mice at this stage showed GFP positivity on flow cytometry, so this is essentially a comparison of early versus late disease (or, with reference to neoadjuvant treatment, disease that would be managed surgically). Percentages of live CD45+ cells are shown in Figure S1. Figure 2A shows an overview of immune cell types detected by flow cytometry, and their expression as a percentage of total CD45+ cells. Late tumors induced significant increases in the proportion of total T cells, CD4+ T cells, regulatory T cells (Tregs), and plasmacytoid dendritic cells. Statistically non-significant increases in NK cells, B cells and conventional dendritic cells were also observed.

**Figure 2.**
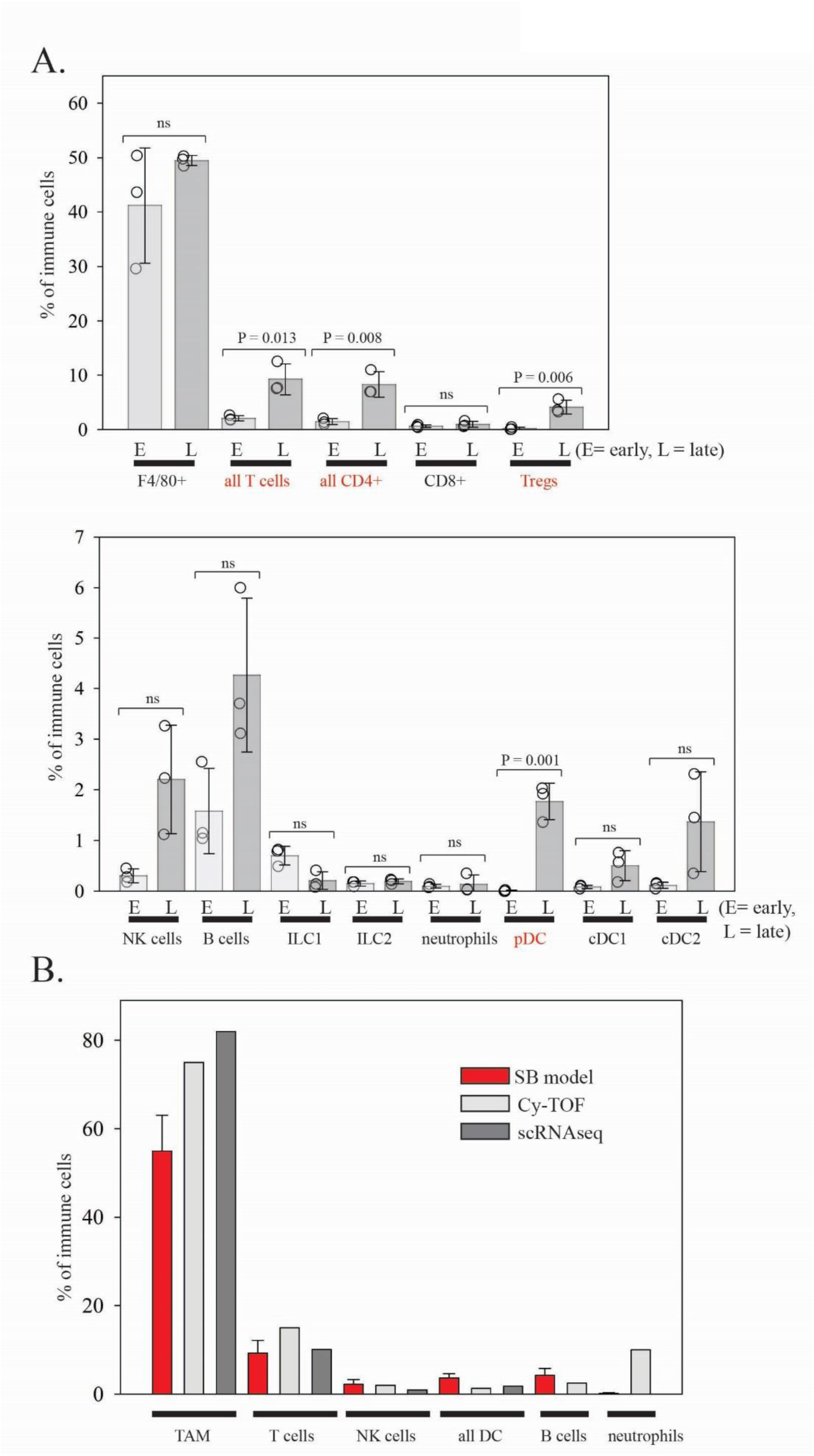
Spectral flow cytometry analysis. A. Mice with tumours detectable by optical imaging (dark gray, labeled L for late stage) are compared to mice that were not showing tumours by optical imaging (light gray, labeled E for early stage) (3 mice per group). The top graph shows data for microglia/macrophages (labeled F4/80+ for the marker used) and T cells. The bottom graph shows data for lower abundance immune cell types. Data are expressed as percent of CD45+ immune cells. Bars show the mean, error bars show standard deviation and individual data points are shown as open circles. P values were determined using Student’s t tests or Mann-Whitney rank sum tests where tests for equal variance failed, with a P value < 0.05 considered significant. B. Pooled data from SB model tumour analyses compared to human data on immune cell frequencies. CyTOF data (21 patients, 4 recurrent) are from Friebel *et al*. (22); scRNAseq data (11 newly-diagnosed patients) are from Abdelfattah *et al*. (23). Mean values are shown for these data sets.

Data from late tumors were pooled and compared to two datasets on human glioblastoma. The first of these was a detailed CYTOF study of sixteen newly diagnosed and four recurrent glioblastoma patients (22); the second was the newly-diagnosed subset of eleven patients from a single cell RNAseq study of immune cell types in glioblastoma (23). There is generally good agreement between the three datasets, with mouse percentages falling in the range reported for the human disease (Figure 2B). One exception is B cells, which are slightly higher in the SB model. Friebel *et al.* showed that a pattern of high microglia/macrophage percentage and low T cell percentage is a feature of glioblastoma, with brain metastases showing the opposite (22). The SB model clearly follows the expected pattern for glioblastoma. The proportion of CD8 T cells was small, at one percent of CD45+ cells (range 0.6-1.6 %) and was not significantly higher than in early disease brain (Figure 2A). This CD8+:CD4+ ration is lower than the average for patients in both the CYTOF and scRNAseq datasets, although there is a wide interpatient variation. Han *et al*. showed that a low CD8+ and high CD4+ cells is a strong independent predictor of poor prognosis in glioblastoma, suggesting that the SB model may model a very aggressive form of glioblastoma (24).

### T cell distribution in the SB model

Immunohistochemistry for CD3 was used to study the distribution of the T cell population that was increased in late stage tumors. T cells were rare in the tumour mass (Figure 3A). Increases in T cells were observed primarily in perivascular regions and the leptomeninges of tumour bearing mice, but were very rare in these regions in normal mouse brain (Figure 3B-G). This is similar to what is observed in a study of glioblastoma patients that looked at T cell neuroanatomical distribution in detail (25). FoxP3 immunohistochemistry showed that Tregs followed the same distribution as the overall CD3+ population, but in lower numbers (Figure 4A-H). Tregs were not detected in normal mouse brain (Figure 4G). Figures 4D and F show Tregs lining a blood vessel emanating from the meninges (Figure 5C and E), occupying the space between the pia mater and the blood vessel wall (*i.e.* the Virchow-Robin space). In the meninges, both the total T cell and Treg populations were most commonly present in the vicinity of leptomeningeal blood vessels (Figures 3E, 4C and E). Flow cytometry showed that the Treg population was 58 ± 1 % positive for the proliferation marker Ki67; non-FoxP3-expressing CD4 cells were similar, at 56 ± 4 % positive.

**Figure 3.**
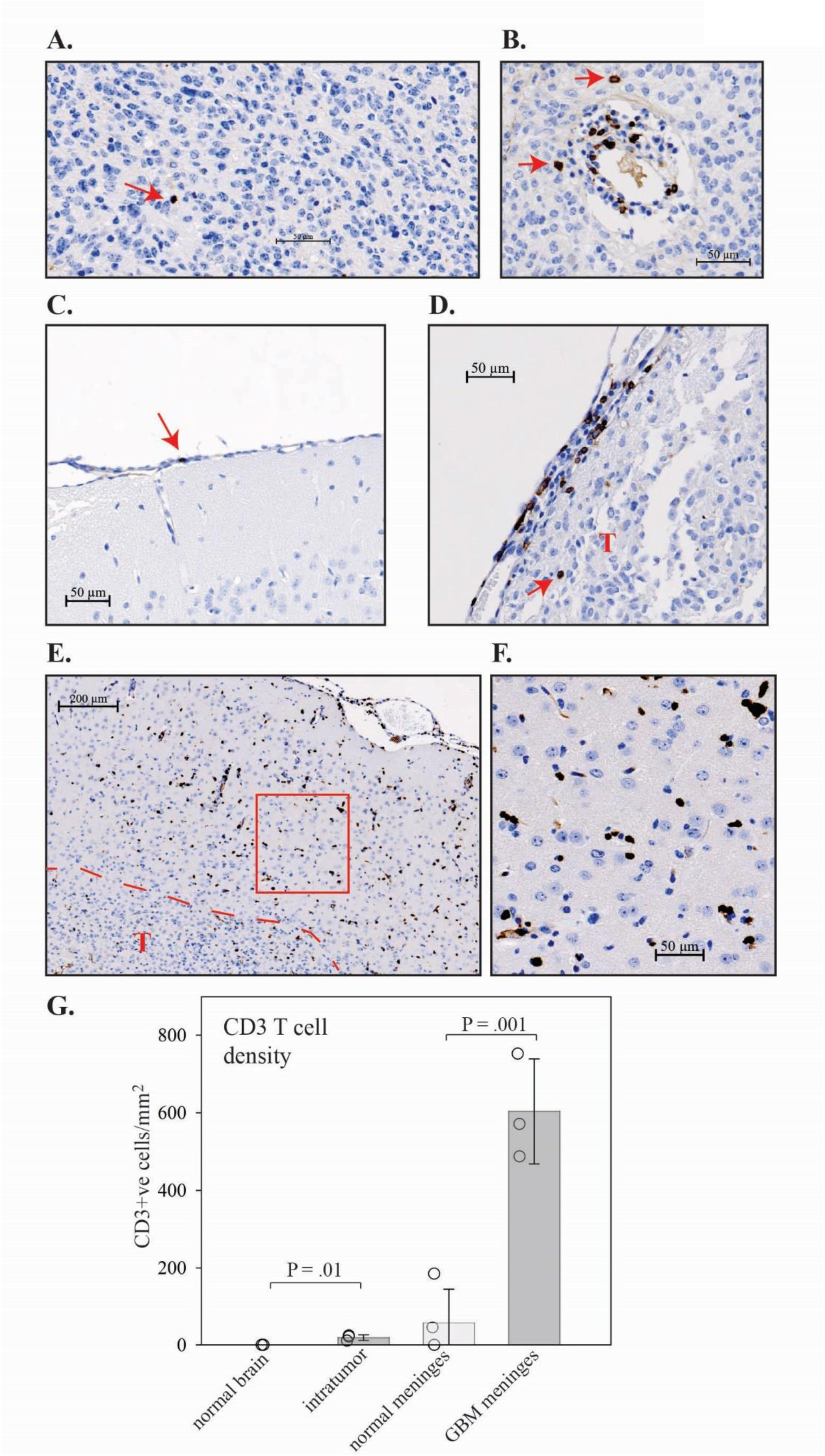
Immunohistochemistry analysis of the overall T cell population. A. Image of tumour parenchyma. CD3+ cells are very sparse, with the single CD3+ cell in this field of view indicated with a red arrow. B. CD3+ cells in a perivascular region. Red arrows show two cells that appear to have exited the perivascular space. C. CD3+ cells are rare in normal mouse leptomeninges. The red arrow indicates one of the four CD3+ cells detected in this whole brain section. D. CD3+ cells are abundant in leptomeningeal regions of tumor-bearing mice. The red T indicates the tumour parenchyma and the red arrow indicates a CD3+ cell in the tumor parenchyma. E. Image showing leptomeninges (top right), brain parenchyma and tumour parenchyma (red dotted line and T). F. Image showing closeup of peritumoral brain parenchyma (area in red from E.) with relatively abundant CD3+ cells relative to the tumour mass. G. Intratumor and leptomeningeal T cell densities were determined for the SB model (three mice) and normal mouse brain.

**Figure 4.**
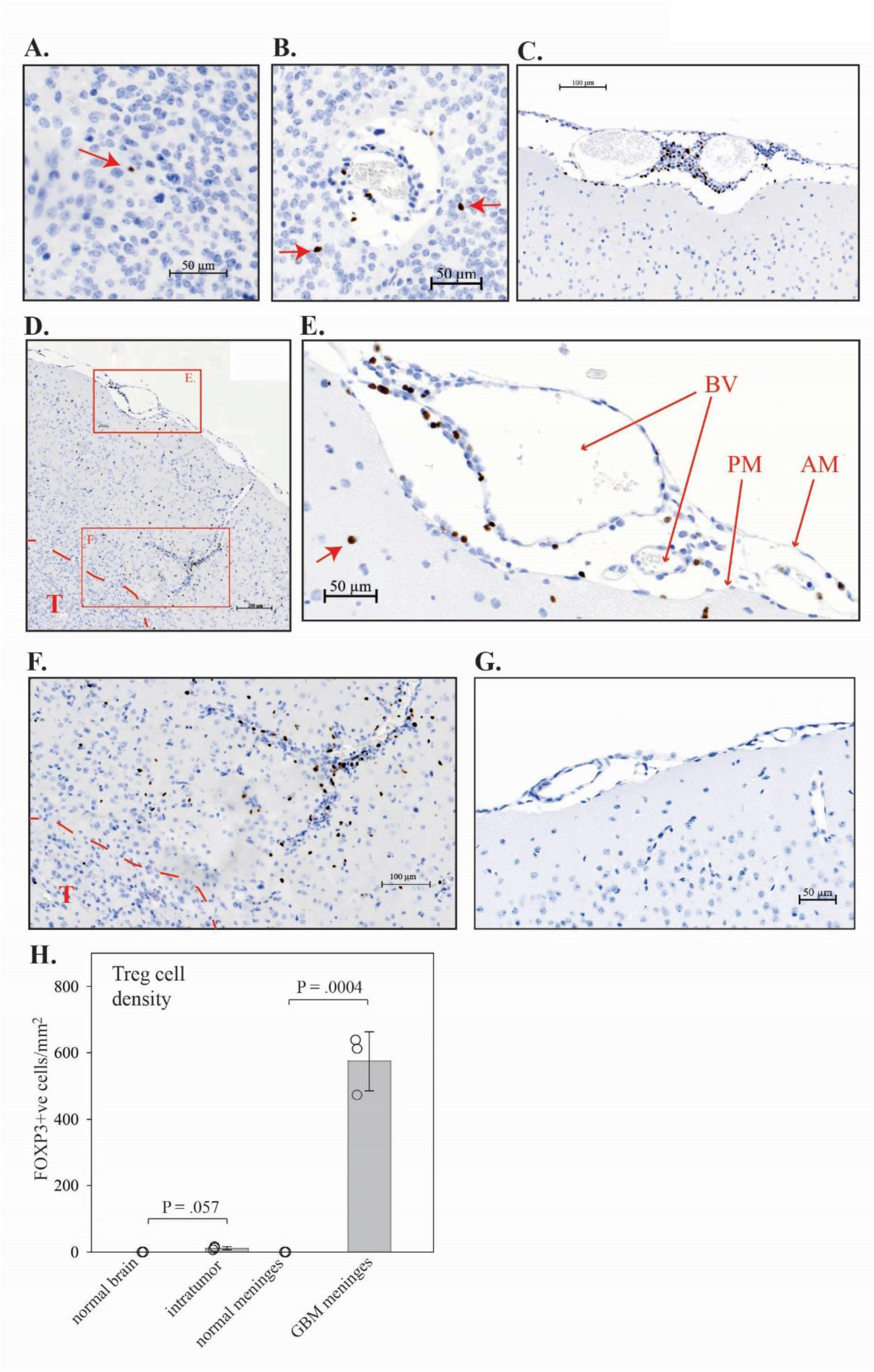
Regulatory T cell characterization. Immunohistochemistry for FoxP3 was performed on normal mouse brain and tumours from the SB model. A. FoxP3+ cells are very sparse in the tumour parenchyma. Red arrow indicates the single FoxP3+ cell present in this field of view. B. FoxP3+ cells in perivascular region. Red arrows indicate two FoxP3+ cells present in the tumour parenchyma, while more FoxP3+ cells are present in the perivascular region. C. Example of FoxP3+ cells in the vicinity of leptomeningeal blood vessels. D. Region showing leptomeninges (red box E), brain parenchyma with blood vessel/pia mater invagination (red box F), and tumour mass (red dotted line and T). E. Close up of leptomeninges. Red arrows with small heads indicate: blood vessels (BV); arachnoid membrane (AM); and pia mater (PM). Red arrow with large head indicates a FoxP3+ cell that is present in brain parenchyma. F. Close up of pia mater invagination/blood vessel with abundant paravascular FoxP3+ cells. G. FoxP3+ cells are not present in normal mouse leptomeninges. H. Intratumor and leptomeningeal Treg densities were determined for the SB model (three mice) and normal mouse brain.

**Figure 5.**
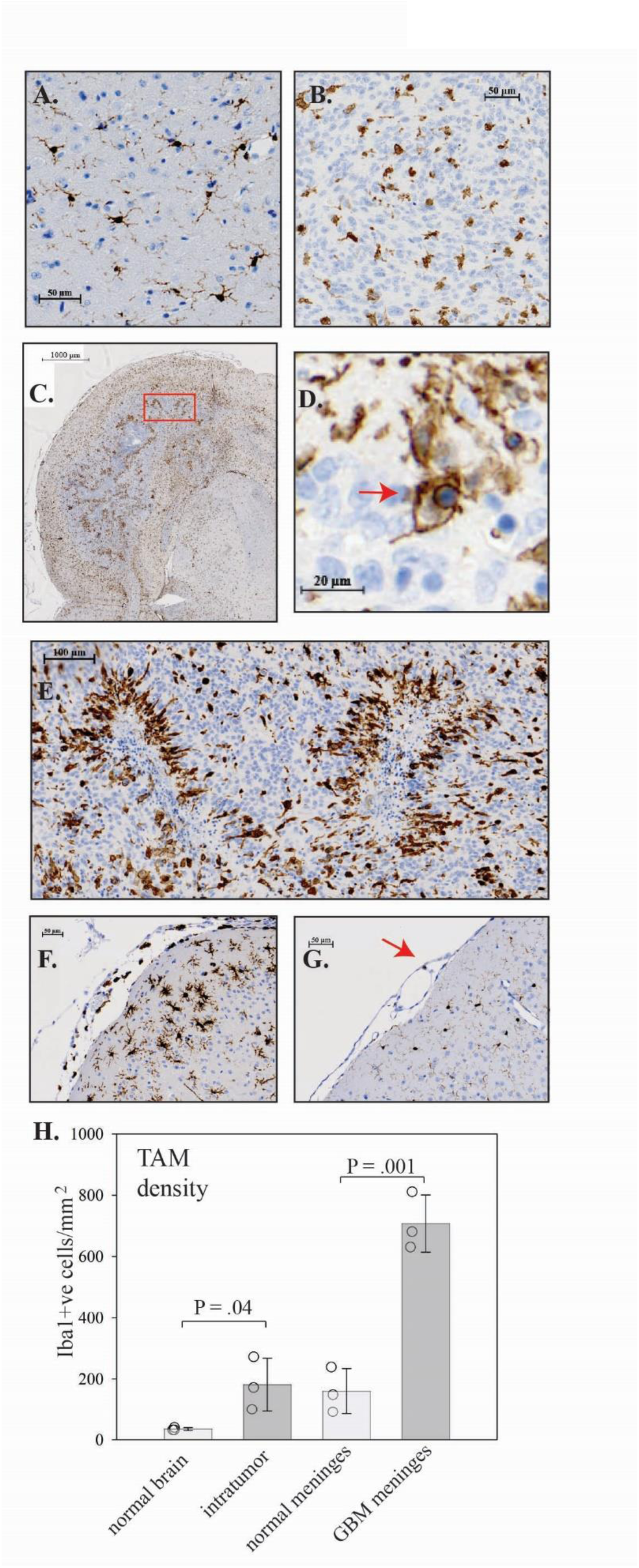
TAM characterization in SB model. Immunohistochemistry for Iba1 was performed on normal brain and tumours from the SB model. A. Iba1 expression in normal mouse brain (for comparison). B. Iba1 expression in a tumor parenchyma without vasculature or necrosis. C. Iba1 expression in in a hemisphere with extensive necrosis. D. Close up of TAM at border of necrotic region showing phagocytic activity (red arrow). E. Recruitment of TAM to necrotic regions. A close up of the region indicated by the red box in C is shown. Hematoxylin staining shows small dark blue apoptotic cells. F. Iba1staining shows abundant macrophages in the leptomeninges of a tumor-bearing mouse. G. Iba1 staining of leptomeninges in normal mouse brain for comparison. Red arrow indicates the single positive cell in this region. H. Quantitation of TAM density in tumor parenchyma without vasculature or necrosis (intratumor) and in meninges in three tumor bearing mice.

### Immune cells in the choroid plexus in the SB model

Border sites for immunosurveillance of the brain include the meninges and perivascular regions, and also the choroid plexus, which has fenestrated capillaries that ultimately connect back to cervical lymph nodes (9). In the SB model, CD3 immunohistochemistry in one mouse showed that T cells (CD3+) were present in the lateral ventricle choroid plexus at higher levels than observed in normal mouse choroid plexus (Figure S2, right top and bottom panels). T cells accumulated at the choroid plexus base, where fibroblasts form a barrier separating the choroid plexus from cerebrospinal fluid (26). This barrier is normally tightly sealed by adherens junctions and tight junctions, but becomes permissive to small molecules and cells with inflammation. The accumulation of cells suggest that the barrier is intact here, so that T cells in the choroid plexus would not be exposed to cerebrospinal fluid. These T cells did not express FoxP3. However, the presence of T cells in choroid plexus was only observed in one of three mice with large tumors that were examined, so engagement of this border site appears to be less frequent than observed for leptomeninges.

### PD-1 and CTLA-4 immune checkpoint expression in the SB model

The expression of PD-1 and CTLA-4 in late-stage tumors, as determined by flow cytometry, is shown in Figure S3A. The CD8 T cell population in mice with detectable tumours was 16 ± 3 % positive for CTLA-4 and 41 ± 15 % positive for PD-1; expression of the latter is restricted to effector memory CD8 T cells (Figure S3B and C). This is consistent with a small subset of cytotoxic T cells that are dysfunctional via an exhaustion mechanism. Non-FoxP3-expressing CD4+ cells were 46 ± 8 % positive for CTLA-4 and 59 ± 13 % positive for PD-1. As expected, Tregs were 95 ± 1 % positive for CTLA-4 and, based on their frequency, are therefore the largest source of CTLA-4 expression in SB model glioblastoma. Based on analysis of scRNAseq data from eleven newly-diagnosed glioblastoma patients (23), Tregs also appear to be a major source of CTLA-4 in human glioblastoma (Figure S3D).

### NK cells

NK cells comprised 1-3% of CD45+ cells, very similar to what has been observed in patients. Levels were consistently higher than seen in non-tumour brain, although this was not significant due to variation in the extent of the increase in tumours (Figure 2A). NK maturation was assessed using the four stage maturation process defined by CD11b and CD27 expression (27). NK cells showed a markedly reduced maturity in comparison to NK cells in the spleen from the same mouse (Figure S4A). Tumour NK cells also had lower expression of KLRG1, another marker of NK cell maturation (Figure S4B). In glioblastoma patient tumours, NK cells are also predominantly in immature states (22).

### Tumor-associated microglia/macrophages (TAM)

Across multiple experiments and multiple tumors (n=5), TAM in tumour tissue, as assessed by the marker F4/80, comprised on average 55% of CD45+ cells (range of 48-67%). This overlaps with the range reported for glioblastoma *IDH*wt patients, although the average for this was higher at approximately 75%. Immunohistochemistry for Iba1 confirmed abundant TAM in non-necrotic tumour regions, with a slightly higher density than observed in normal mouse brain (Figure 5A, B and H). As described in human glioblastoma, TAM adopted an ameboid morphology, in contrast to the ramified morphology observed in normal brain (28). TAM clustered around regions of necrosis in the tumours, where they were observed to be engulfing dying cells, also as seen in the human disease (29)(Figure 5C-E). TAM were markedly increased in the leptomeninges of tumor-bearing mice compared to normal mice (Figure 5F, G and H), as has been observed in glioblastoma patients (25).

### TGFB1 expression

Immunohistochemistry for TGFB1 showed that it is expressed at high levels in perinecrotic regions (Figure 6A). The close up image of a necrotic region shown in Figure 6B (left panel) shows that TGFB1 expression rims a region of apoptotic cells (small dark blue spots with hematoxylin stain) and follows the same distribution as TAM surrounding the apoptotic cells, but not TAM in the surrounding tumor mass, as shown with Iba1 immunohistochemistry on an adjacent section (Figure 6B, right panel); this is consistent with TAM undergoing reprogramming to an immunosuppressive state at these sites as a consequence of their efferocytosis activity. TGFB1 was also present in the leptomeninges at higher levels than in normal mouse (Figure 6C). There is also a diffuse increase in signal throughout the tumor-bearing brain in comparison with normal brain stained in parallel (Figure 6C). A subset of strongly positive-staining cells in the leptomeninges are likely to be Tregs, which, in addition to requiring TGFB1 for their induction, also express TGFB1 bound to the cell surface protein GARP (30).

**Figure 6.**
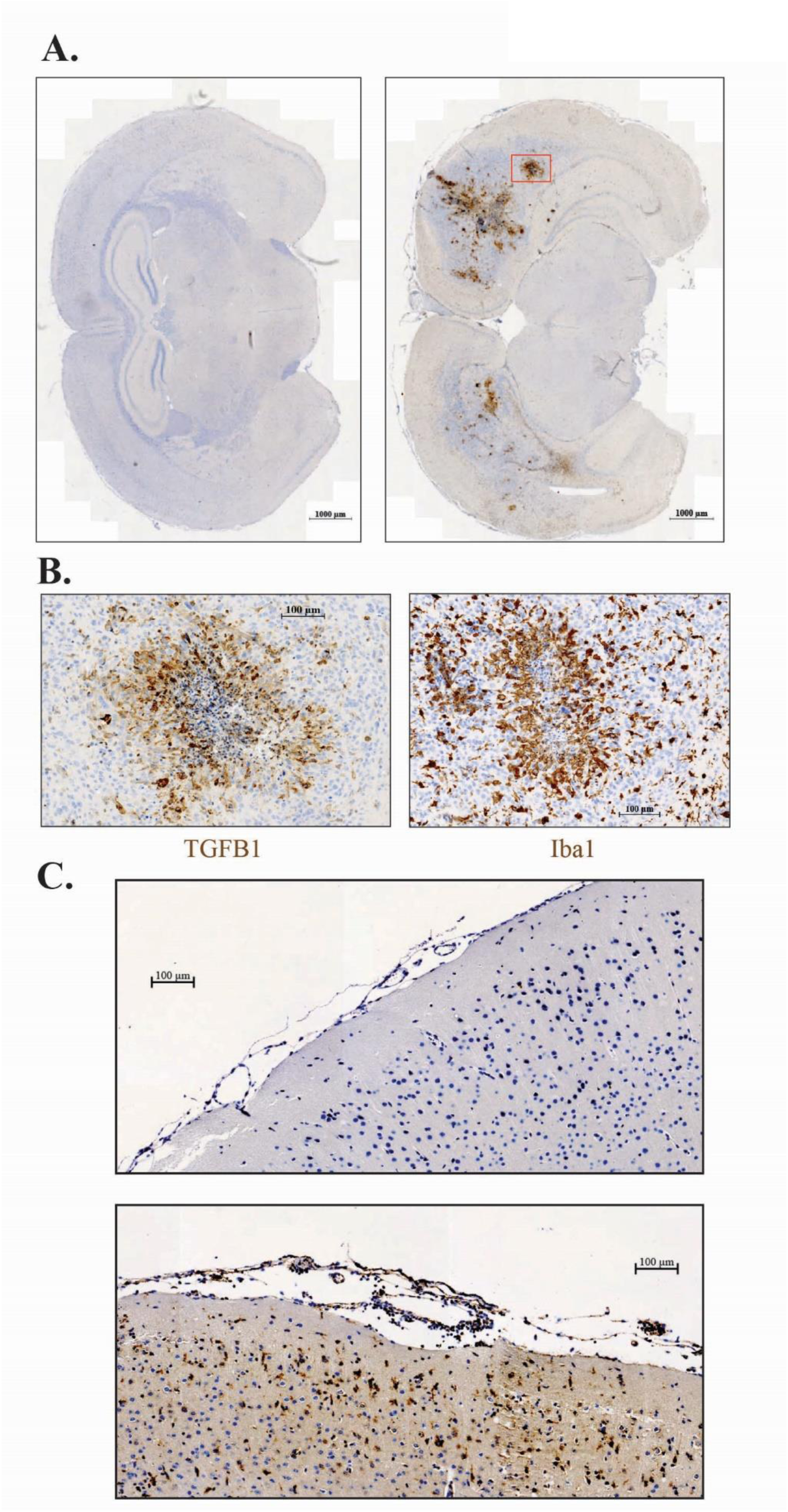
TGFB1 expression in SB model. A. The left panel shows a whole brain coronal section from a normal FVB/N mouse stained for TGFB1 expression; the right panel show the same for an SB model tumor-bearing mouse. B. The left panel shows a close up of the necrotic region indicated by the red box in A. The right panel shows the same region in an adjacent section stained for Iba1. C. Close up of leptomeninges and brain parenchyma in a normal FVB/N mouse (left panel) and an SB model mouse (right panel) stained for TGFB1. Images were taken at matched higher contrast than used in A to emphasize the diffuse expression of TGFB1 in brain parenchyma.

## DISCUSSION

Given the success of immunotherapy in other cancer types, there is intense interest in determining whether immunotherapies such as immune checkpoint inhibition can work in glioblastoma. Having mouse models that accurately model the human disease are an important aspect of this research. The *de novo* glioblastoma model described by Wiesner *et al*. was described over fifteen years ago, but has not been widely adopted, probably because of the challenges of performing intracerebroventricular injections accurately and reproducibly in neonatal mice. Here we show that the use of an adaptor for a standard mouse stereotaxic apparatus improves this aspect of the model significantly. As shown previously, the model is versatile, both because it can be adapted to use different oncogenic drivers and because it can be used across different mouse strains (11, 13, 31). In terms of using this model for preclinical immunotherapy evaluation, Genoud *et al*. previously isolated a cell line from this model and showed that this had a mutational load similar to that reported for human glioblastoma (16). They also showed that injection of this cell line into mice gave tumours that were unresponsive to immune checkpoint inhibition, similar to what is observed in patients. This supports the use of the orthotopic version of this model for translational studies. Here we have performed a detailed immune cell analysis of the original autochthonous version of this model, using spectral flow cytometry and immunohistochemistry. This shows that the model closely resembles the human disease immunologically. Overall proportions of recruited immune cells are similar to what is observed in patients. The increases in T cells in the meninges and perivascular regions, the inhibition of NK cell maturation, and the behaviour of microglia/macrophages in tumor tissue and necrotic regions are all similar to what has been reported for patients (22, 25, 29).

As mentioned in the introduction, the concept of immunoprivilege in the brain is undergoing revision in the scientific literature, with the growing recognition that border sites immunosurveil the brain via the cerebrospinal fluid (9). Very early studies showing that transplants of mouse tumor tissue into rat brain survived if they were wholly surrounded by brain parenchyma, but were rejected if transplanted close to the ventricles, showing that immunoprivilege was location sensitive (32). Large glioblastoma tumors confront the problem that they are more likely to engage border site immunosurveillance once they occupy a larger region of the brain. We show here that large tumors in the SB model do breach immunoprivilege, inducing a marked increase in T cells. T cell accumulation is particularly pronounced at leptomeningeal/perivascular border sites. These T cells are 40-60% positive for PD-1 and therefore may be affected by PD-1 targeted immune checkpoint inhibition. Based on recent literature, these T cells probably come from cranial bone, enter the dura mater via microchannels and then cross into the subarachnoid space at arachnoid cuff exit (ACE) points (33, 34). ACE points occur where blood vessels cross the arachnoid barrier, leaving a cell-permeable space around the vessel (9, 34). In the SB model, leptomeningeal T cells are most abundant in the vicinity of meningeal blood vessels, consistent with this route of entry into the subarachnoid space, although further studies would be needed to confirm this.

This immune response is countered by immunosuppressive Tregs, which make up approximately half of the CD4 T cell population in the SB model. Notably, Tregs are potent inhibitors of dendritic cells (35, 36). The Tregs are present in perivascular spaces where the initial event in brain immunosurveillance, mixing of brain interstital fluid with CSF, takes place (37); they are therefore positioned to block adaptive immune responses at their inception. Induced Tregs are generated from CD4 T cells by TCR engagement with class II MHC on dendritic cells in the presence of TGFβ (38, 39). Along with an increase in immunosuppressive Tregs, NK cells in the SB model are predominantly in immature states which have low cytotoxic activity. The repression of NK cell maturation is also mediated by TGFβ (40, 41). Key mechanisms for innate (NK cell) and adaptive (T cell) immunity are therefore both repressed in the SB model by mechanisms that are known to depend on TGFβ.

The SB model accurately reproduces the palisading necrosis that is a defining characteristic of human glioblastoma. Originally thought to be simply a consequence of tumors outgrowing their blood supply, these necrotic regions are now known to be important drivers of glioblastoma malignancy (42). They are generated by thrombotic events in the tumor vasculature initiated by overexpression of pro-coagulant factors in the tumor microenvironment. Necrotic regions generate large numbers of apoptotic cells, a behaviour that can be reproduced in cell culture by oxygen deprivation of glioblastoma patient-derived organoids (29). Caspase cleavage of membrane transporters causes the membranes of apoptotic cells to become leaky; as a consequence of this they have a distinct small molecule secretome that is highly chemoattractive to macrophages (43). Microglia/macrophages are aggressively recruited to clear away apoptotic cells before they go on to immunogenic secondary necrosis, a process known as efferocytosis. In addition to clearing away cells, efferocytic macrophages are reprogrammed to express immunosuppressive molecules such as TGFβ (44–48). Efferocytosis is an essential component of inflammation resolution that is initiated by neutrophil apoptosis. Glioblastomas appear to co-opt this process for immune evasion (29, 49). This is evident in the SB model where necrotic regions generate large numbers of apoptotic cells and recruit macrophages. Macrophages in the vicinity of apoptotic cells express high levels of TGFβ, while macrophages in other regions do not, consistent with reprogramming of macrophages by efferocytosis in necrotic regions. Whole sections of mouse brain illustrate the magnitude of this process in glioblastoma, with apoptotic cell generation likely far exceeding what occurs in normal tissues. While there are undoubtedly other sources of TGFβ, immunohistochemistry suggests that necrotic regions are a major source in large tumors in the SB model. The dark TGFβ staining around necrotic regions likely is due to intracellular newly synthesized TGFβ and latent TGFβ associated with extracellular matrix. Compared to normal tissue stained in parallel, glioblastoma sections show a weak brown staining overall suggesting active soluble TGFβ throughout the brain, including the leptomeninges. Necrotic region-generated TGFβ potentially impacts multiple aspects of the immune system, including the generation of Tregs from CD4 T cells and the suppression of NK cell maturation described above. This immunosuppressive role for necrotic regions is consistent with recent spatial transcriptomics studies on patient samples, which showed a high immunosuppression signature in perinecrotic regions (50).

As discussed in the Introduction, a clinical trial of the PD-1 inhibitor pembrolizumab showed pharmacodynamic evidence for activity but limited or no overall clinical benefit (6, 8). The findings here provide potential explanations for this, with large tumors breaching the brain “immunoprivilege” state and initiating an immune response, but adapting to this by inducing efferocytosis to drive the expression of immunosuppressive TGFβ. Strategies to repress TGFβ signaling, either directly or indirectly via efferocytosis inhibition, should be tested in combination with immune checkpoint inhibition. The modified SB model described here is a rapid and clinically-relevant glioblastoma model for preclinical evaluation of these strategies. Limitations of studies on glioblastoma patient tissue are that border site tissue is often absent and necrotic regions are often excluded when selecting tumor tissue for analysis. These are key regions that need to be studied in depth for a comprehensive understanding of glioblastoma immunology. A significant limitation of this study is that cranial bone and dura mater are not included in our analyses; these are also key regions with respect to immune responses to glioblastoma (33) and future studies on these regions in the SB model would be valuable.

## Supporting information

Lui et al 2026 manuscript

## ETHICS

All mouse experiments were done under a protocol approved by the University of Ottawa Animal Care Committee (OHRIe-4326).

## FUNDING

IAJL: Canadian Institutes of Health Research Project Grant #162180 (co-applicant); Brain Cancer Canada Discovery Grant.

MA: Canadian Institutes of Health Research Project Grant; Terry Fox Research Institute Program Project Grant

## CONFLICT OF INTEREST

MA reported grants from Dragonfly Therapeutics and Actym Therapeutics outside the submitted work. None declared for ML, OJM, MLW, TM, JW, MA, IAJL.

## AUTHORSHIP

Experimental design: ML, OJM, MA, IAJL

Experimental implementation and/or analysis: ML, OJM, MLW, TM, JW, MA, IAJL

Interpretation of data: ML, OJM, JW, MA, IAJL

Writing of the manuscript: IAJL wrote the initial draft manuscript, ML, OJM, MLW, TM, JW and MA reviewed the draft and all authors have read and approved the final manuscript.

## DATA AVAILABLITY

Data from this study are freely available for non-commercial purposes upon request to the corresponding author.

## ACKNOWLEDGEMENTS

We gratefully acknowledge Histology/Imaging/Staining services provided by the Louise Pelletier Core Facility (RRID: SCR_021737) at the Department of Pathology and Laboratory Services, Faculty of Medicine, University of Ottawa.

