## Supplementary material for "Characterization of tumour interactions with the immune system in an autochthonous mouse model of glioblastoma": Lui et al 2026 manuscript

### SUPPLEMENTARY METHODS AND FIGURES

**Neonatal Mouse Intraventricular Stereotactic Injection Protocol for SB Model**
(based on Wiesner *et al*.(1) and Calinescu *et al*.(2))

1. Prepare equal volumes of DNA and in vivo jetPEI transfection mix to the desired total volume for the number of mice to be injected according to manufacturer's protocol. An injection for one mouse contains a total of 1.0 ug DNA in 2.0 uL of injection volume.
   - Using 4-plasmid model with PT2/C-Luc//PGK-SB13, PT3.5/CMV-EGFRvIII, PT2/shp53/GFP4, PT/Caggs-NRASV12, DNA amount is 1:1:1:1.
   - jetPEI N/P ratio used is 7.
2. Grab a good amount of bedding from the nest in the home cage. Place some bedding in a cut-open glove on top of an ice slurry to maintain home cage scent on pups and reserve the rest for use at the stereotactic frame and recovery heating pad.
3. Collect three neonatal mice from the cage and induce anesthesia by placing on top of the glove and bedding on the ice slurry. Monitor color and pinch reflex until extremities pale and no reflex occurs.
   - Never leave the home cage with no pups remaining with the mother! Pace the surgery so that the first group of pups are recovered and ready to be returned to the cage when the last group of pups can be prepared for anesthesia.
4. While waiting for anesthesia induction, prepare surgery stage at the stereotactic frame. Mount a 10-uL Hamilton syringe with a 1-inch 30G needle to the injector and load with the transfection mix. Secure and level neonatal mouse head adaptor on the ear and bite bars and place a small ice pack with bedding on top underneath the adaptor.
5. When pups are anesthetized, quickly move one pup over to the stereotactic frame and cradle head in the adaptor with straight alignment.
6. Clean the head with a small amount of ethanol on a cotton swab. Make an incision in the scalp to expose the skull.
7. Using the needle, determine the coordinates for lambda. Move needle +1.5 AP and +0.7 ML to the left ventricle.
8. From the depth of the surface of the skull, insert needle -1.5 DV and begin injection at 0.7 uL/min for 2.0 uL.
9. After the injection is finished, slowly withdraw the needle and use tissue glue to close the incision.
10. Move the pup to a heated recovery pad with some bedding and monitor for return of color and movement.
11. Repeat anesthesia and injection until all neonatal mice in the cage have been injected.
    - A new group of pups can be taken from the cage and placed on ice when halfway through the prior group's injections. Remember to never leave the mother without pups.
12. At the end of surgery, provide cereal and extra cotton squares for the mother to minimize overgrooming of pups. Note whether the mother interacts with the returned pups appropriately. Bring the cage back to the housing room and perform wellness checks on the cage at 4 and 24 hours post surgery.

**Spectral Flow Cytometry Panel**

| Marker | Clone | Company |
|---|---|---|
| CD11b BUV661 | M1/70 | BD Biosciences |
| CD69 BUV737 | H1.2F3 | BD Biosciences |
| CD27 Brilliant Violet 510 | LG.3A10 | BD Biosciences |
| KLRG1 Brilliant Violet 605 | 2F1 | BD Biosciences |
| NKp46 Brilliant Violet 650 | 29A1.4 | Biolegend |
| PD-L1 Brilliant Violet 711 | 10F.9G2 | Biolegend |
| PD-1 Brilliant Violet 750 | 29F.1A12 | Biolegend |
| Ki67 Brilliant Violet 786 | B56 | BD Biosciences |
| CD62L BB515 | MEL-14 | BD Biosciences |
| CD49a BB700 | Ha31/8 BD | Biosciences |
| EOMES PerCP-eFluor710 | Dan11mag | eBioscience |
| ST2 PE | U29-93 BD | Biosciences |
| CD45 PE/Dazzle 594 | 30-F11 | Biolegend |
| CD3 PE-Cyanine 5 | 145-2C11 | Biolegend |
| CD19 PE-Cyanine 5 | 6D5 | Biolegend |
| F4/80 PE-Cyanine 5 | BM8 | Biolegend |
| GATA3 PE-Cyanine 7 | L50-823 | BD Biosciences |
| DX5 Alexa Fluor 647 | DX5 | Biolegend |
| DNAM-1 APC/Fire 750 | 10E5 | Biolegend |
| CD8 BUV395 | 53-6.7 | BD Biosciences |
| CD4 BUV661 | GK1.5 | BD Biosciences |
| KLRG1 Brilliant Violet 421 | 2F1 | BD Biosciences |
| CD80 Brilliant Violet 510 | 16-10A1 | BD Biosciences |
| TIGIT Brilliant Violet 605 | 1G9 | BD Biosciences |
| CD44 Brilliant Violet 650 | IM7 | BD Biosciences |
| T-bet Brilliant Violet 711 | 4B10 | Biolegend |
| CD3 BB700 | 500A2 | BD Biosciences |
| EGR2 PE-Cyanine 7 | erongr2 | eBioscience |
| TOX APC | REA473 | Miltenyi Biotech |
| F4/80 Alexa Fluor 647 | BM8 | Biolegend |
| FOXP3 Alexa Fluor 700 | MF-14 | Biolegend |
| CD11c BUV395 | HL3 | BD Biosciences |
| CD103 BUV563 | M290 | BD Biosciences |
| CCR7 BUV805 | 4B12 | BD Biosciences |
| NKp46 Brilliant Violet 421 | 29A1.4 | Biolegend |
| CD172a Brilliant Violet 605 | P84 | BD Biosciences |
| MHC-II Brilliant Violet 650 | M5/114.15.2 | Biolegend |
| Ly6C Brilliant Violet 785 | 1A8 | Biolegend |
| CD24 PerCP-Cy5.5 | M1/69 | Biolegend |
| Siglec F PerCP-eFluor710 | 1RNM44N | eBioscience |
| CD68 PE | FA/11 | BD Biosciences |
| Ly6G PE-Cyanine 7 | D7 | BD Biosciences |
| CD317 Alexa Fluor 700 | 927 | Biolegend |
| KLRG1 BUV563 | 2F1 | BD Biosciences |
| CD25 Brilliant Violet 605 | PC61 | BD Biosciences |
| FOXP3 eFluor 450 | FJK-16s | eBioscience |
| CTLA-4 PE-Fire 810 | UC10-4B9 | Biolegend |

*Figure S1. CD45 positive immune cells in the SB model.* Spectral flow cytometry was performed as described in Figure 2. Top panels show three mice with early disease, bottom panels show three mice with late disease (tumors visible by optical imaging). The percent of live CD45 positive cells is shown in the top left corner of each plot.

*Figure S2. Immune cells in the choroid plexus.* Staining for CD3, FOXP3 and Iba1 are shown for choroid plexus from normal mouse and a SB model mouse with a late stage tumor. The bottom panels show the base of the choroid plexus where it attaches to the ventricle wall.

*Figure S3. Immune checkpoint expression.* T cell populations were analyzed for expression immune checkpoint proteins by spectral flow cytometry. A. PD-1 and CTLA4 expression by regulatory T cells (FoxP3+), CD4 cells (CD4+/FoxP3-), CD8 cells (CD8+). Data are from tumours from three mice. Bars show the mean, error bars show standard deviation and individual data points are shown as open circles. B. Distribution of naïve, central memory and effector memory CD8+ T cells from matched spleen and tumour. C. Left and middle panels show CD69 and PD-1 expression in naïve, central memory and effector memory CD8+ T cells in spleen and tumor from the same mouse. Right panel shows PD-1 expression in ILC2 cells in spleen and tumor from the same mouse. D. CTLA4 and PD-1 mRNA expression in glioblastoma patient immune cell types. scRNAseq data (11 newly-diagnosed patients) are from Abdelfattah *et al.* (23).

*Figure S4. NK cell characterization in the SB model.* A. Spectral flow cytometry data was used to characterize the maturation state of NK cells. R0 and R1 are functionally immature states, while R2 and R3 are more functionally mature states. Matched spleen and tumour from three  are mice are compared. Percentages in R0, R1, R2 and R3 states are indicated. Bar graph shows pooled data for CD11-ve and CD11+ve spleen and tumor NK cells. * indicates a P value < .05 by two-tailed t tests. B. Expression of the NK cell maturation marker KLRG1 on matched spleen and tumour NK cells from three mice.

Figure S1.

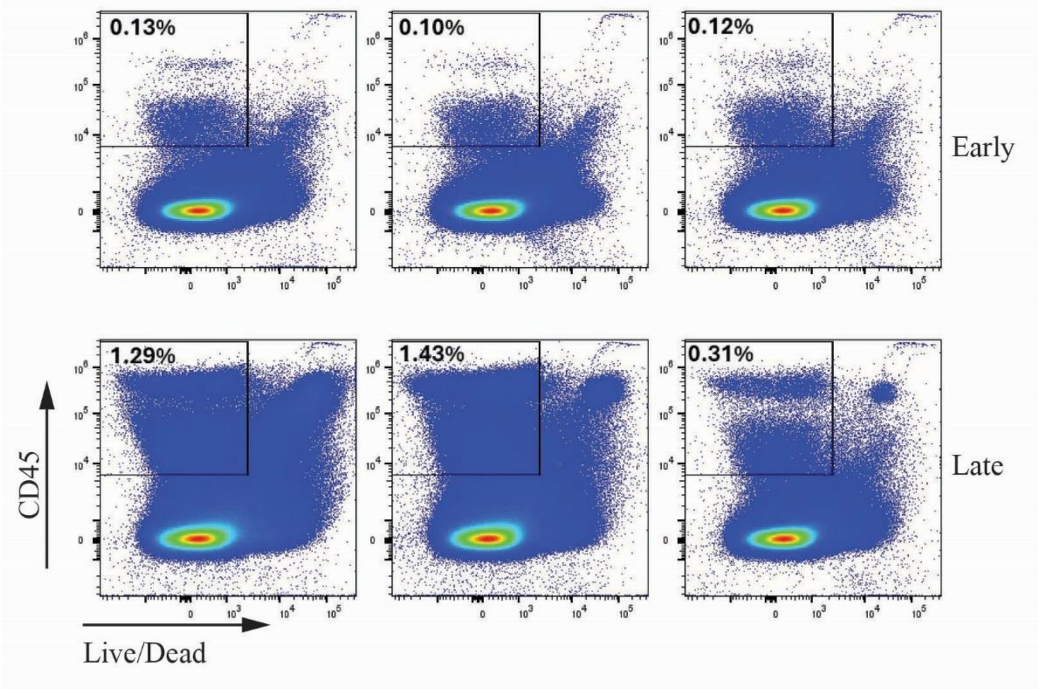

Figure S2.

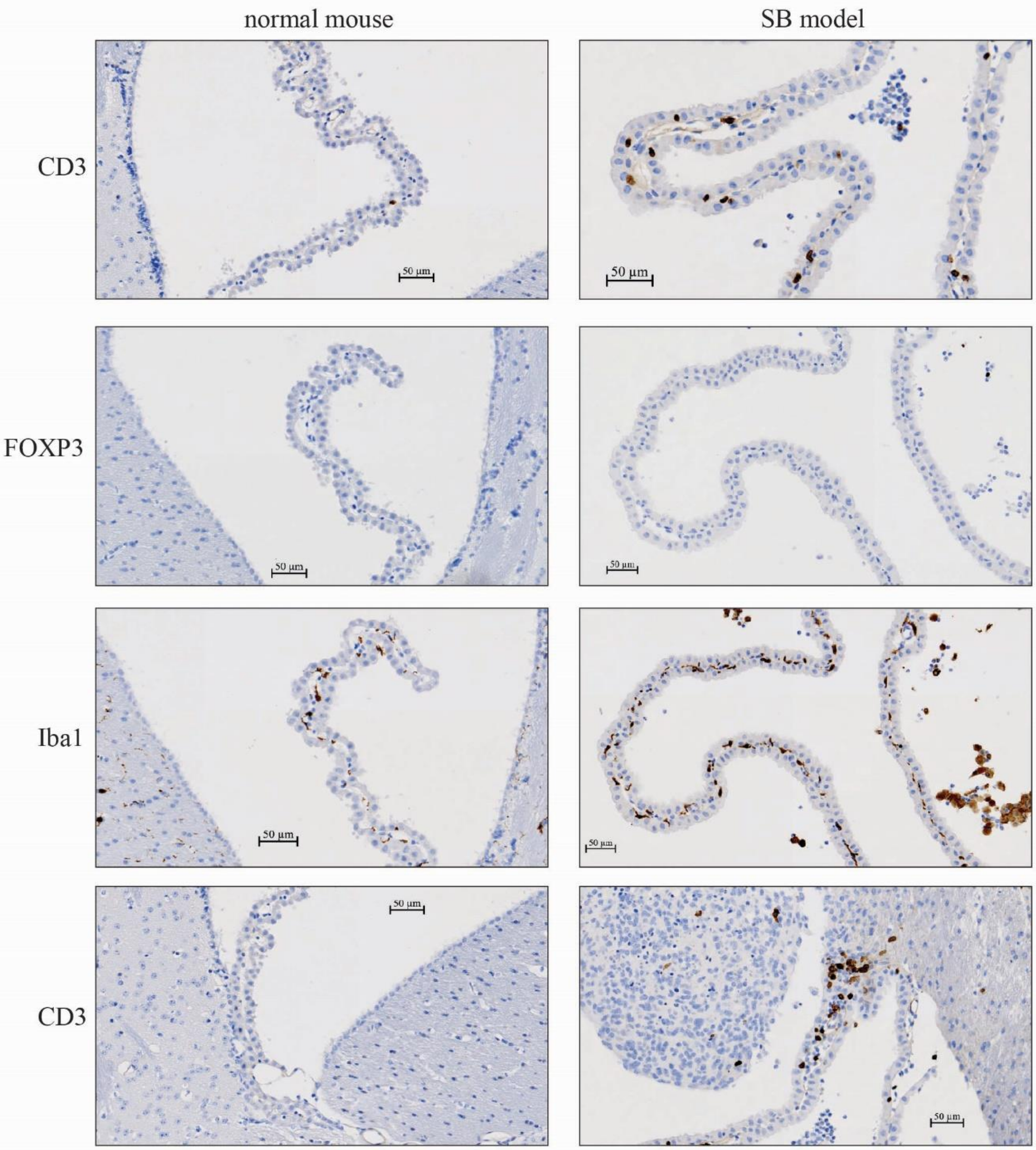

Figure S3.

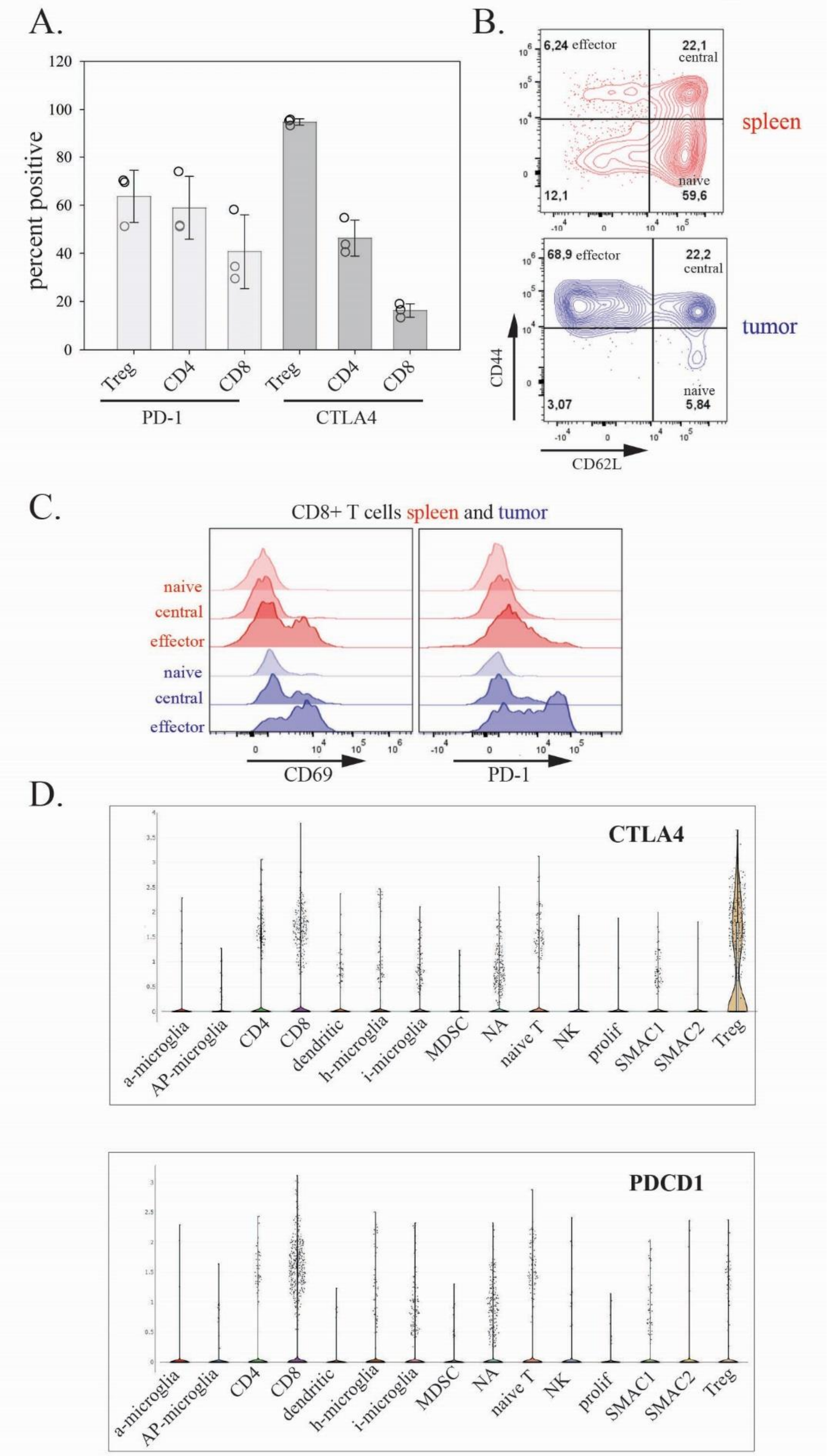
A.
percent positive
120
100
80
60
40
20
0
Treg
CD4
CD8
Treg
CD4
CD8
PD-1
CTLA4
B.
6,24 effector
22,1 central
spleen
12,1
naive 59,6
68,9 effector
22,2 central
tumor
3,07
naive 5,84
CD44
CD62L
C.
CD8+ T cells spleen and tumor
naive
central
effector
naive
central
effector
CD69
PD-1
D.
CTLA4
a-microglia
AP-microglia
CD4
CD8
dendritic
h-microglia
i-microglia
MDSC
NA
naive T
NK
prolif
SMAC1
SMAC2
Treg
PDCD1
a-microglia
AP-microglia
CD4
CD8
dendritic
h-microglia
i-microglia
MDSC
NA
naive T
NK
prolif
SMAC1
SMAC2
Treg

Figure S4.

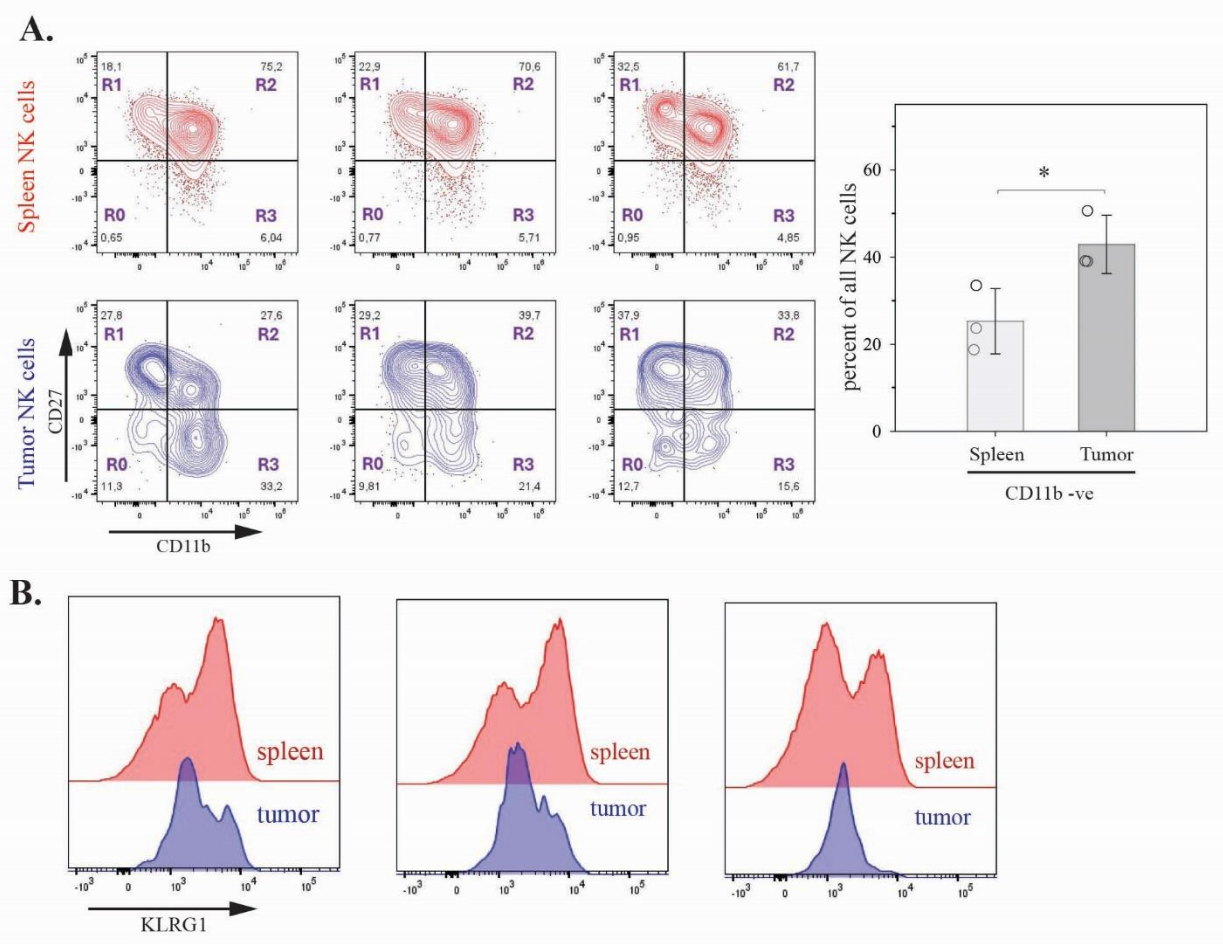
A.
Spleen NK cells
Tumor NK cells
R1
R2
R0
R3
18,1
75,2
0,65
6,04
22,9
70,6
0,77
5,71
32,5
61,7
0,95
4,85
27,8
27,6
11,3
33,2
29,2
39,7
9,81
21,4
37,9
33,8
12,7
15,6
CD27
CD11b
percent of all NK cells
*
Spleen
Tumor
CD11b -ve
B.
spleen
tumor
KLRG1
